# Distinct effects of disease-associated TREM2 R47H/+ and T66M mutations on iPSC-derived microglia

**DOI:** 10.1101/2022.10.05.511003

**Authors:** Jay Penney, William T Ralvenius, Anjanet Loon, Oyku Cerit, Vishnu Dileep, Blerta Milo, Hannah Woolf, Li-Huei Tsai

**Affiliations:** Picower Institute for Learning and Memory, Massachusetts Institute of Technology, Cambridge, MA, 02139, USA; Department of Brain and Cognitive Sciences, Massachusetts Institute of Technology, Cambridge, MA, 02139, USA; Broad Institute of MIT and Harvard, Cambridge, MA, USA

## Abstract

Genetic findings have highlighted key roles for microglia in the pathology of neurodegenerative conditions such as Alzheimer’s disease (AD). Distinct mutations in the microglial protein TREM2 (triggering receptor expressed on myeloid cells 2) are associated with different forms of neurodegeneration in humans; R47H/+ mutations increase AD risk, while loss-of-function mutations such as TREM2 T66M result in more severe forms of neurodegeneration. We employed gene editing and stem cell models to gain insight into the effects of these mutations on human iPSC-derived microglia. We found divergent effects of TREM2 R47H/+ and T66M mutations on gene expression, with R47H/+ cells exhibiting a pro-inflammatory gene expression signature. Both the TREM2 R47H/+ and T66M mutations caused similar impairments in microglial movement and the uptake of multiple substrates, while R47H/+ microglia were hyper-responsive to inflammatory stimuli, consistent with their gene expression signature. We developed an *in vitro* laser-induced injury model in neuron-microglia co-cultures, finding an impaired injury response by TREM2 R47H/+ microglia. Furthermore, in xenotransplantation experiments, mouse brains transplanted with TREM2 R47H/+ microglia exhibited reduced synaptic density. Consistently, we observed upregulation of multiple complement cascade components in TREM2 R47H/+ microglia, suggesting that inappropriate synaptic pruning may underlie the effect. Thus, these findings identify shared and distinct effects of these two TREM2 mutations on microglial gene expression and function. While the TREM2 T66M mutation impairs microglial movement and uptake processes, the TREM2 R47H/+ mutation additionally confers multiple potentially detrimental effects on human microglia, likely to underlie its association with AD.

## Introduction

Alzheimer’s disease (AD) is a devastating neurodegenerative condition with no cure (Canter et al., 2016). The AD brain is characterized by widespread neurodegeneration, neuroinflammation, and the presence of hallmark protein aggregates composed primarily of amyloid-β (Aβ) or hyperphosphorylated tau protein (amyloid plaques and neurofibrillary tangles, respectively; De Strooper and Karran, 2016). Familial forms of AD (fAD) are well established to result from dominantly inherited mutations that affect Aβ production and/or processing by neurons, thus neuronal mechanisms have historically been the focus of AD research (Hardy and Higgins, 1992; Makin, 2018). More recently, however, large-scale genome-wide association studies (GWAS), have broadened our view of AD pathogenesis by identifying many non-neuronal genetic risk factors for sporadic forms of AD (sAD; Bellenguez et al., 2022; Escott-Price and Hardy, 2022; Kunkle et al., 2019; Lambert et al., 2013). Remarkably, almost half of the >90 prioritized sAD risk genes identified thus far are expressed more highly in microglia than in any other brain cell type, indicating that microglial dysfunction can be an active driver in the disease process (Cahoy et al., 2008; Escott-Price and Hardy, 2022; Lewcock et al., 2020; Penney et al., 2020). Among these microglia-specific sAD risk factors is the subject of this study, the triggering receptor expressed on myeloid cells 2 (TREM2) (Guerreiro et al., 2013; Jonsson et al., 2013).

Microglia perform numerous important functions in the developing and adult brain. As innate immune cells, microglia continually survey their environment and can phagocytose components of dead cells, protein aggregates and other debris, while also regulating inflammation via cytokine release in response to various stimuli (Li and Barres, 2018). Additionally, microglia can more directly impact neurodevelopment by providing trophic support and sculpting neural circuits via synaptic pruning (Salter and Stevens, 2017). In contrast, while microglial activities are essential for proper neuronal development and function, chronic inflammation due to microglial activation is a hallmark of neurodegenerative disease that can directly incite neuronal damage (Lewcock et al., 2020). Synaptic pruning by microglia can also be inappropriately re-activated, contributing to synapse loss during the neurodegenerative process (Hammond et al., 2019).

In 2013, heterozygous point mutations in TREM2 were identified as risk factors for AD, most notably the TREM2 R47H/+-causing mutation which increases AD risk by roughly three-fold (Guerreiro et al., 2012; Jonsson et al., 2013; Zhou et al., 2019). More than a decade earlier, however, homozygous deletion mutations in TREM2 and its binding partner TYROBP were identified as causative mutations for polycystic lipomembranous osteodysplasia with sclerosing leukoencephalopathy (PLOSL), an aggressive early onset disease characterized by bone cysts, demyelination, neurodegeneration, and dementia (Errichiello et al., 2019; Paloneva et al., 2002). Additional homozygous loss-of-function mutations, including TREM2 T66M, which prevents TREM2 glycosylation and thus cell surface expression, were subsequently shown to cause PLOSL and a frontotemporal dementia (FTD)-like syndrome (Dardiotis et al., 2017; Kleinberger et al., 2014). Cell surface expressed TREM2 can also be cleaved, liberating extracellular soluble TREM2 (sTREM2) species, and the TREM2 intracellular domain (ICD) (Kleinberger et al., 2014; Thornton et al., 2017).

TREM2 has been shown to regulate many microglial processes. TREM2 knockout mice exhibit impaired developmental synaptic pruning and impaired sensing and clearance of debris following myelin injury (Filipello et al., 2018; Nugent et al., 2020; Poliani et al., 2015). In AD mouse models, TREM2 knockout impairs the ability of microglia to sense and respond to amyloid plaques, also resulting in detrimental effects in tau models (Gratuze et al., 2020; Keren-Shaul et al., 2017; Lee et al., 2021; Leyns et al., 2017; Wang et al., 2016, 2015). Inflammatory responses by microglia in amyloid and tau models are generally dampened by loss of TREM2 (Griciuc et al., 2019; Jay et al., 2017; Leyns et al., 2017). Studies examining TREM2 R47H mutations have often reported similar effects to knockout mutations, with R47H microglia in amyloid models exhibiting impaired plaque localization and response (Cheng-Hathaway et al., 2018; Song et al., 2018). The apparent effects of the R47H mutation in tau mouse models have varied, with reports of increased or decreased tau seeding, microglial inflammation and synaptic/cognitive effects (Gratuze et al., 2020; Leyns et al., 2019; Sayed et al., 2021).

Many microglial risk genes for AD, including TREM2, are relatively poorly conserved between mouse and human, suggesting that the use of human induced pluripotent stem cell (iPSC) models to understand their function is of particular importance (Penney et al., 2020). Accordingly, iPSC-based studies have also examined the effects of TREM2 null, R47H/+ and T66M mutations. Each mutations has often, but not always, been reported to reduce uptake of brain-relevant substrates such as Aβ (Andreone et al., 2020; Liu et al., 2020; McQuade et al., 2020; Piers et al., 2020; Reich et al., 2021). Metabolic impairments have also been observed in microglia carrying these mutations, while reported effects on movement and chemotaxis have been mixed (Jairaman et al., 2022; Piers et al., 2020; Reich et al., 2021). Normal or impaired inflammatory responses in TREM2 null and T66M iPSC-microglia have mostly been reported, while different studies have shown either increased or decreased inflammation in cells carrying the R47H mutation (Andreone et al., 2020; Brownjohn et al., 2018; Cosker et al., 2021; Garcia-Reitboeck et al., 2018; Hall-Roberts et al., 2020; Liu et al., 2020; Piers et al., 2020). Similarly, single-nucleus and bulk RNA sequencing experiments from AD patients carrying R47H/+ mutations have observed a pro-inflammatory gene expression signature, though not in all cases (Korvatska et al., 2020; Sayed et al., 2021; Yingyue Zhou et al., 2020).

Here, we use gene editing and iPSC differentiation to compare the effects of isogenic TREM2 R47H/+ and T66M mutations in human microglia. We find a shared set of genes commonly affected by the two TREM2 mutations, as well as gene expression changes specific to each mutation, highlighted by a pro-inflammatory signature in R47H/+ microglia. In functional assays, we find similar effects of R47H/+ and T66M mutations on the uptake of multiple substrates and on cellular movement. Consistent their inflammatory gene signature, we find that TREM2 R47H/+ microglia exhibit exaggerated cytokine release and gene expression changes in response to inflammatory stimuli. We also find impaired responses of R47H/+ microglia in a novel laser-induced neuronal injury model. Further, in xenotransplantation experiments in mice, we find that synaptic density is reduced specifically by TREM2 R47H/+ iPSC-microglia. Thus, we identify numerous functional and gene expression alterations in human microglia resulting from TREM2 of R47H/+ and T66M mutations, including multiple effects specific to the R47H/+ mutation that may underlie its association with AD.

## Results

### TREM2 R47H/+ and T66M mutations differentially affect microglial gene expression

Beginning with an iPSC line derived from a healthy 75-year-old female, we performed CRISPR/Cas9-mediated gene editing (see Methods; Cong et al., 2013) to introduce point mutations corresponding to either the R47H or T66M substitutions in TREM2 protein (Figure 1A). Following screening of putative gene edited iPSC clones, we moved forward with two clones carrying heterozygous R47H-causing mutations (R47H/+) and two clones carrying homozygous T66M-causing mutations, associated with AD and PLOSL, respectively (Figure 1B). An additional clone from the CRISPR screening that remained un-edited at the TREM2 locus, as well as the parental iPSC line, were used as isogenic controls. We next used a modified protocol from McQuade et al. to differentiate these six iPSC lines into iPSC-derived microglia cells for further validation and analysis (see Methods; McQuade et al., 2018).

**Figure 1.**
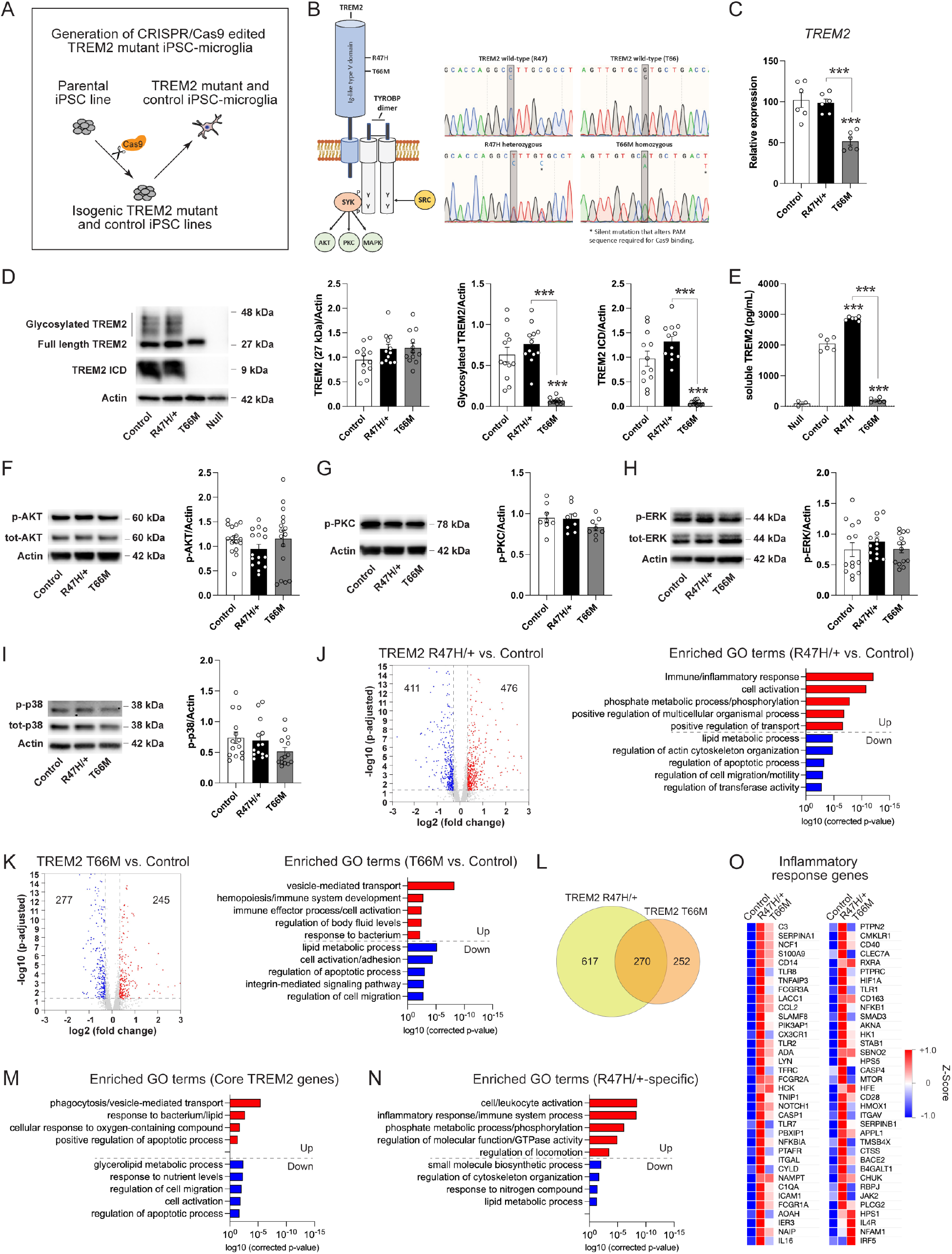
TREM2 R47H/+ and T66M mutations commonly alter a core set of genes with R47H/+ microglia also exhibiting a pro-inflammatory signature. (A) Schematic of CRISPR mutagenesis and generation of TREM2 mutant microglia. (B) Diagram of TREM2 protein and downstream effectors. Locations of the R47H and T66M mutations are indicated. Chromatogram traces showing successful editing of the R47H and T66M-coding sequences in TREM2. (C) Quantification of *TREM2* mRNA level in TREM2 mutant and isogenic control microglia. n=6 for each (3 samples from clone A and 3 from clone B). ***p<0.001. 1-way ANOVA with Tukey’s test. (D) Images and quantification of western blots for different forms of TREM2. Actin serves as loading control. n=12 for all. ***p<0.001. 1-way ANOVA with Tukey’s test. (E) Quantification of ELISA for soluble TREM2 levels in culture media from control and mutant microglia. n=6 for all. ***p<0.001. 1-way ANOVA with Tukey’s test. Blot and quantification from control and mutant microglia of (F) phospho-AKT levels, n=16 for all, (G) phospho-PKC levels, n=8 for all, (H) phospho-ERK levels, n=14 for all, and (I) phospho-p38 levels. n=14 for all. (J) Volcano plot and enriched GO terms for genes differentially expressed between control and TREM2 R47H/+ microglia. (K) Volcano plot and enriched GO terms for genes differentially expressed between control and TREM2 T66M/+ microglia. (L) Venn diagram of the overlapping and unique genes differentially expressed in R47H/+ and T66M microglia. Enriched GO terms for (M) ‘core’ genes commonly affected by R47H/+ and T66M mutations and (N) genes uniquely affected by the R47H/+ mutation. (O) Heatmap of the gene expression Z-Score for genes in the ‘Inflammatory response’ GO term in control and TREM2 mutant microglia that showed increased expression in one or both of TREM2 R47H/+ and T66M microglia.

Upon iPSC-microglia maturation, we first examined *TREM2* mRNA expression in these cells. While we found similar expression of *TREM2* transcript in R47H/+ and control cells, we observed a significant decrease in *TREM2* expression in cells carrying T66M mutations, potentially indicating destabilization of the transcript by this mutation (Figure 1C). For this and all future experiments, unless otherwise noted, an equal number of samples from clone A and clone B of each genotype were analyzed and the results pooled.

We next used western blotting to examine whether TREM2 protein levels and glycosylation/maturation were affected by the R47H/+ and T66M mutations. Lysates from control and R47H/+ cells show a main band at 27 kDa, corresponding to full length TREM2 protein, as well as higher molecular weight species corresponding to glycosylated forms of TREM2 (Figure 1D). As expected, glycosylated forms of TREM2 were not observed in lysates from T66M microglia, while levels of glycosylated TREM2 were similar in control and R47H/+ cells (Figure 1D; Kleinberger et al., 2014). Likewise, similar levels of the TREM2 intracellular domain (ICD) were observed in samples from control and R47H/+ microglia, while TREM2 ICD was absent from T66M lysates (Figure 1D). Lysates from TREM2 null cells, included as a control for antibody specificity, showed no signal for any TREM2 species (Figure 1D). We then used ELISA to quantify sTREM2 levels in media from control, R47H/+ and T66M microglial cultures (Figure 1E). We observed an increase in sTREM2 produced from R47H/+ cells, relative to controls, and almost undetectable levels of sTREM2 in T66M microglia media (Figure 1E). Elevated sTREM2 levels have also been reported in R47H/+ AD patients (Piccio et al., 2016).

We then examined whether cell signaling pathways engaged downstream of TREM2 signaling were altered by TREM2 R47H/+ or T66M mutations. We found no significant differences in the levels of phosphorylated forms of Ak strain transforming (AKT), protein kinase C (PKC), or the mitogen-activated protein kinases (MAPKs) extracellular signal-regulated kinase (ERK) or p38 in microglia carrying either TREM2 mutation vs. controls (Figures 1F-1I). These results indicate that the major signaling modules downstream of TREM2 are not appreciably affected by the R47H/+ or T66M mutations in un-stimulated microglia.

We next sought to characterize gene expression changes that result from these TREM2 mutations in iPSC-microglia. To that end, we performed RNA sequencing on mRNA from control, R47H/+ and T66M microglia (see Methods). We identified 887 genes as differentially expressed (>0.3 log2 fold change, p-adjusted<0.05) in R47H/+ microglia compared to controls, and 522 genes in T66M microglia vs. controls (Figure 1J and 1K). Gene ontology analysis identified inflammatory responses and cell activation as the most enriched biological processes among upregulated genes in R47H/+ cells (Figure 1J). Vesicle-mediated transport-related genes were the most enriched among those upregulated by the T66M mutation, with transport process also being enriched among R47H/+ upregulated genes (Figure 1J and 1K). Analysis of downregulated genes from R47H/+ and T66M cells revealed a shared enrichment for biological processes affecting lipid metabolism, apoptotic processes and cell migration (Figure 1J and 1K).

Next, we examined overlap among the genes we found differentially expressed in R47H/+ and T66M microglia. We found a core set of 270 genes that whose expression was significantly altered, in the same direction, by both the R47H/+ and T66M mutations (Figure 1L). These shared differentially expressed genes accounted for more than half of all genes whose expression was significantly affected by the T66M mutation (Figure 1L). In contrast, we found that more than 600 genes were uniquely differentially expressed in microglia carrying the TREM2 R47H/+ mutation (Figure 1L).

We found that upregulated ‘core’ TREM2 genes (increased by both R47H/+ and T66M mutations) were most enriched for phagocytosis and vesicle-mediated transport pathways (Figure 1M). Biological processes that were enriched among core downregulated genes mirrored those found in our initial comparisons between R47H/+ and T66M vs. control microglia, affecting processes such as glycerolipid metabolism, cell migration and apoptotic processes (Figure 1M). We found significant enrichment for only a handful of biological processes among T66M-specific genes: cytoskeleton and organelle localization, vesicle-mediated transport, and regulation of fibrinolysis among upregulated genes, and lipid metabolic processes in downregulated genes. In contrast, genes specifically altered by the TREM2 R47H/+ mutation showed strong enrichment for a number of biological processes, by far the most prominent being related to cell activation and inflammation (Figure 1N). Examination of the expression levels of genes from the biological process ‘inflammatory response’ that were upregulated by one or both of the R47H/+ or T66M mutations underscores the pro-inflammatory gene expression signature of TREM2 R47H/+ microglia (Figure 1O).

### Microglial movement and cellular uptake are impaired by TREM2 R47H/+ and T66M mutations

To determine whether the gene expression changes we observed in R47H/+ and T66M microglia resulted in altered cellular functions, we next sought to perform a series of *in vitro* functional assays on these cells. We first examined the ability of control, R47H/+ and T66M microglia to take up a number of brain-relevant substrates. TREM2 has been shown to play a role in developmental synaptic pruning in the mouse brain, and microglia have also been demonstrated to engage in inappropriate synaptic pruning in adult Alzheimer’s disease mice (Filipello et al., 2018; Hong et al., 2016; Salter and Stevens, 2017). We purified synaptosomes (isolated preparations of synaptic membrane) from wild-type mouse brains via differential sucrose centrifugation (see Methods), then added 20 μg/mL of synaptosomes to microglial monocultures. After 3 hours, we fixed and immuno-stained the cultures with antibodies recognizing the synaptic protein synaptophysin (SVP38), as well as the microglia marker IBA1 (Figure 2A). We observed synaptophysin signal inside control, and to a lesser extent R47H/+ and T66M microglia, indicating synaptosomes were internalized by these cells (Figure 2A). To more accurately quantify synaptosome uptake by control and TREM2 mutant microglia, we next conjugated our synaptosome preparations to the pH sensitive dye pHrodo-green, which fluoresces more strongly in low pH cellular compartments such as lysosomes (see Methods). pHrodo-labeled synaptosomes were incubated with our microglia for 3 hours, then the cells were collected, washed, and subjected to flow cytometry to quantify synaptosome uptake (Figure 2D). We found that pHrodo-synaptosome signal in both R47H/+ and T66M microglia was reduced relative to controls, indicating impaired synaptosome uptake by microglia carrying TREM2 mutations (Figure 2D).

**Figure 2.**
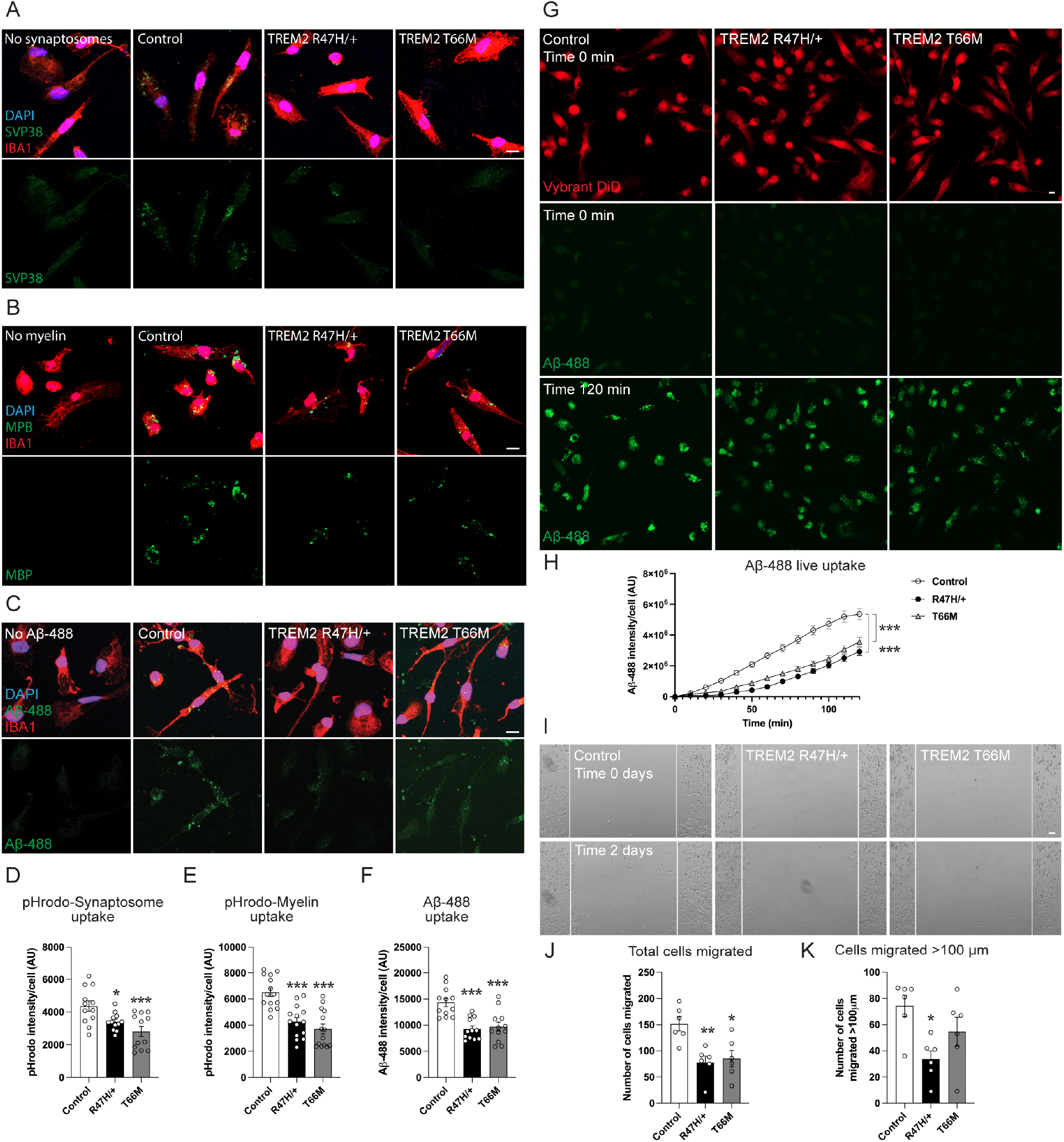
TREM2 R47H/+ and T66M mutations impair uptake processes and movement in microglia. (A) Images of TREM2 mutant and isogenic control microglia stained for IBA1 (red) and SVP38 (green) following 3 hours uptake of mouse synaptosomes. A no-synaptosome condition using control cells is included to verify SVP38 specificity. Scale bar = 10 μm. (B) Images of TREM2 mutant and control microglia stained for IBA1 (red) and MBP (green) following 3 hours uptake of mouse myelin. A no myelin condition using control cells is included to verify MBP specificity. Scale bar = 10 μm. (C) Images of TREM2 mutant and control microglia stained for IBA1 (red) following 3 hours uptake of Aβ_42_-488 (green). A no-Aβ_42_-488 condition using control cells is included to verify signal specificity. Scale bar = 10 μm. (D) Quantification of pHrodo-synaptosome signal in flow cytometry experiments using TREM2 mutant and control microglia. n=12 for all. *p<0.05, ***p<0.001. 1-way ANOVA with Tukey’s test. (E) Quantification of pHrodo-myelin signal in flow cytometry experiments using TREM2 mutant and control microglia. n=14 for all. ***p<0.001. 1-way ANOVA with Tukey’s test. (F) Quantification of Aβ_42_-488 signal in flow cytometry experiments using TREM2 mutant and control microglia. n=12 for all. ***p<0.001. 1-way ANOVA with Tukey’s test. (G) Live imaging of microglial Aβ_42_-488 (green) uptake. Microglia are labeled with Vybrant DiD (red). Scale bar = 10 μm. (H) Quantification of Aβ_42_-488 uptake over the 2-hour time series. n>18 for all, from clone B cells, representative of 2 independent experiments, one using each clone ***p<0.001. 1-way ANOVA with Tukey’s test. (I) Images of scratch assay, immediately following scratch wound (top) and after 2 days of re-population (bottom). Scale bar = 50 μm. (J) Quantification of the total number of cells that migrated into the scratch area. n=6 scratch experiments for all. *p<0.05, **p<0.01. 1-way ANOVA with Tukey’s test. (K) Quantification of the number of cells that migrated >100 μm into the scratch area. N=6 scratch experiments for all. *p<0.05. 1-way ANOVA with Tukey’s test.

Microglia also help to clear damaged myelin from the brain, a process that in mice is impaired by TREM2 loss-of-function mutations (Nugent et al., 2020; Poliani et al., 2015). Using differential centrifugation, we isolated the myelin fraction from wild-type mice (see Methods). Incubation of microglial cultures with 40 ug/mL of myelin for 3 hours, followed by immuno-staining for IBA1 and the myelin protein MBP, revealed robust myelin uptake in control microglia, with qualitatively less MBP signal observed in R47H/+ and T66M microglia (Figure 2B). Using pHrodo-green conjugation and flow cytometry, we quantified myelin uptake in our microglia lines, finding significant reductions in pHrodo-myelin signal in R47H/+ and T66M microglia compared to controls (Figure 2E). Thus, uptake of both synaptosomes and myelin debris is impaired by disease-associated TREM2 mutations.

Aβ is also taken up readily by microglia *in vitro* and *in vivo*, thus we next examined the uptake of 488-labeled Aβ_42_ (Aβ-488) by our iPSC-microglia. Following 3 hours incubation with 200 ng/mL Aβ-488, we fixed and immuno-stained microglial cultures to examine IBA1 and Aβ-488 signal (Figure 2C). As with synaptosomes and myelin, we observed that Aβ-488 signal co-localized with IBA1 microglia, with qualitatively less signal in TREM2 mutant cells. Flow cytometry quantification of Aβ-488 signal in control and mutant microglia confirmed significantly reduced uptake of Aβ-488 by R47H/+ and T66M microglia (Figure 2F). Next, to track Aβ uptake over time, we performed live imaging experiments with our microglia lines (Figure 2G and 2H). We first labeled our microglial cells with the cell permeant dye Vybrant DiD to allow live tracking. Then, following addition of Aβ-488 to the culture media, we collected confocal images of the microglia every 10 minutes for 2 hours. The initially weakly diffuse Aβ-488 signal in the media gradually co-localized with Vybrant DiD-labeled microglia (Figure 2G). Quantification of the fluorescent signal intensity per cell revealed significantly reduced Aβ-488 signal from R47H/+ and T66M cells compared to controls as early as 20 minutes after Aβ-488 addition that persisted throughout the course of the experiment (Figure 2G and 2H). Thus, Aβ-488 uptake is also impaired by both the TREM2 R47H/+ and T66M mutations.

One of the GO terms we found enriched in genes downregulated by both the R47H/+ and T66M mutations alike, was ‘regulation of cell migration’ (Figure 1I and 1J). Thus, we examined cellular movement in control, R47H/+ and T66M microglia using a scratch assay (Figure 2I-2K). Cells were plated at 400k cells/well in 6-well plates. The next day, cells were physically removed (scratched) from the middle portion of the well, and the media was changed to remove floating cells. Then movement back into the scratch area was monitored over time. Two days after the initial scratch wound, the number of cells within a field of view that had migrated back into the scratch area was quantified, demonstrating reduced re-population of the area by R47H/+ and T66M microglia (Figure 2J). We also quantified the number of cells that had traveled >100μm into the scratch area, again finding a reduced number of migrated R47H/+ cells compared to controls (Figure 2K). Together, these findings demonstrate that the TREM2 R47H/+ and T66M mutations cause similar impairments in cellular migration and uptake of multiple physiologically relevant substrates in microglia.

### TREM2 R47H/+ microglia have a pro-inflammatory phenotype

We saw a pro-inflammatory gene expression signature in TREM2 R47H/+ microglia in our RNA sequencing experiments, however, previous human, mouse and iPSC studies have reported conflicting effects of R47H mutations on inflammation (Cosker et al., 2021; Liu et al., 2020; Piers et al., 2020; Sayed et al., 2021; Yongji Zhou et al., 2020) Thus, we next explored whether R47H/+ and T66M iPSC-microglia exhibited altered inflammatory responses. We first treated microglia with the pro-inflammatory bacterial component lipopolysaccharide (LPS, 100 ng/mL) for 2 days, then collected culture media and performed ELISA to examine levels of the pro-inflammatory cytokine interleukin 6 (IL6). LPS treatment increased IL6 levels by at least 8-fold in media from microglia of each genotype, with media from R47H/+ microglia containing significantly more IL6 than either control or T66M microglia media (Figure 3A). We next examined the effects of the cytokine interferon γ (IFNγ, 50 ng/mL) on IL6 production by control and mutant microglia. IFNγ treatment for 2 days also increased levels of IL6 in media from control and TREM2 mutant cells (>3-fold for all), with TREM2 R47H/+ cells again secreting significantly more IL6 than controls (Figure 3A). We then examined the effects of LPS and IFNγ on TNFα production by our microglia. Two-day LPS treatment resulted in at least a 6-fold increased TNFα levels in media from all microglia genotypes, with only R47H/+ microglia showing a statistically significant TNFα increase, which was also qualitatively larger (∼13-fold) than in control (∼8-fold) or T66M (∼6-fold) cells (Figure 3B). Two-day IFNγ treatment increased media TNFα levels only ∼2-fold, without clear differences between control, R47H/+ and T66M cells (Figure 3B). Thus, our observations support the idea that the R47H/+ mutation confers a pro-inflammatory effect on microglia while cells carrying the T66M mutation respond more similarly to control cells.

**Figure 3.**
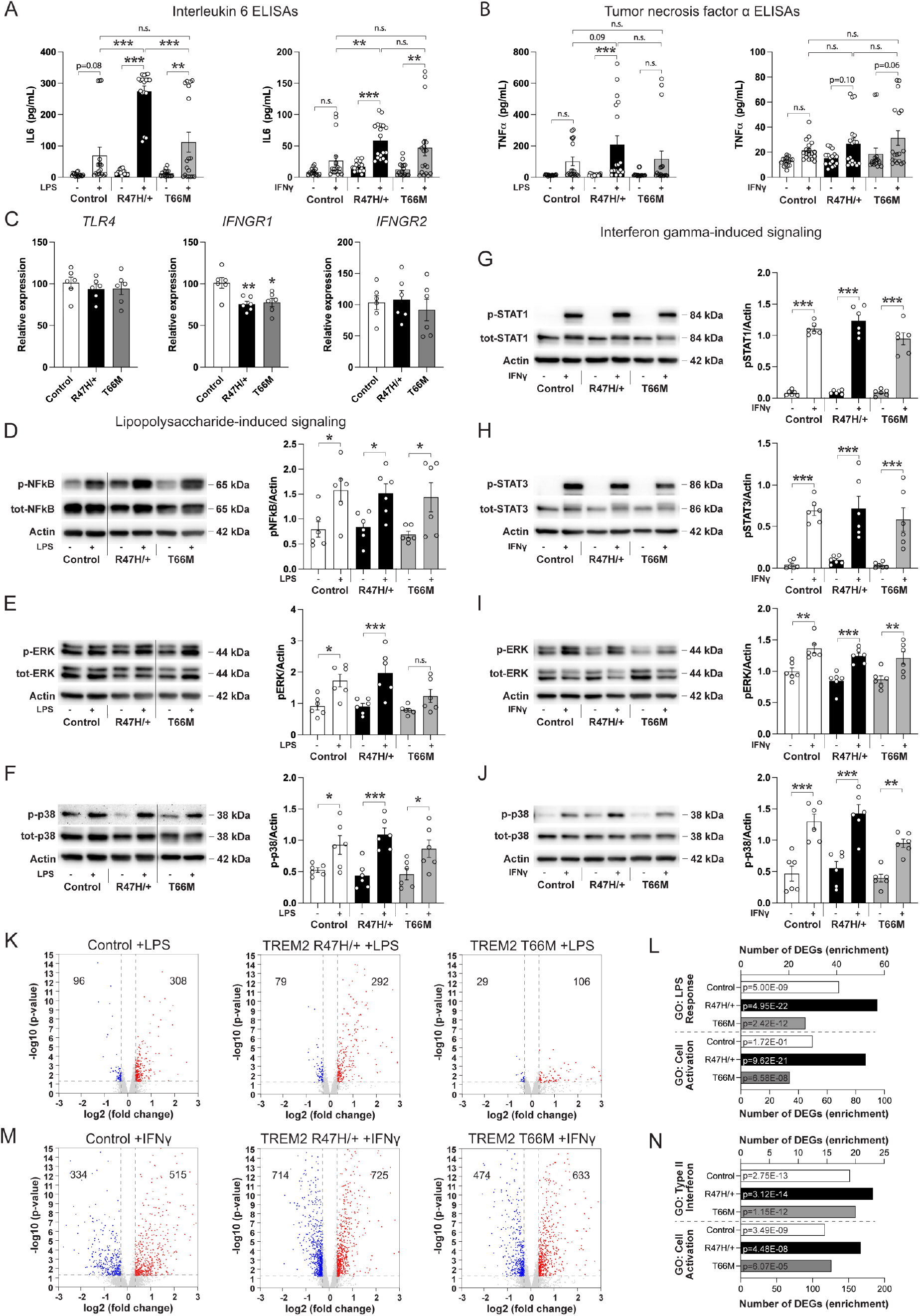
Cytokine release and gene expression changes induced by inflammatory challenge are exaggerated in TREM2 R47H/+ microglia. (A) Quantification of IL6 ELISAs from culture media following treatment with 100 ng/mL LPS (left) or 50 ng/mL IFNγ. n=18 for all. **p<0.01, ***p<0.001. 1-way ANOVA with Sidak’s test. (B) Quantification of TNFα ELISAs from culture media following treatment with 100 ng/mL LPS (left) or 50 ng/mL IFNγ. n=18 for all. ***p<0.001. 1-way ANOVA with Sidak’s test. (C) Quantification of *TLR4, IFNGR1* and *IFNGR2* mRNA levels in TREM2 mutant and isogenic control microglia. n=6 for each. *p<0.05, **p<0.01. 1-way ANOVA with Tukey’s test. Blots and quantifications from LPS-treated control and mutant microglia of (D) phospho-NFkB levels, (E) p-ERK levels and (F) p-p38 levels. n=6 for all. *p<0.05, ***p<0.001. 1-way ANOVA with Sidak’s test. Blots and quantifications from IFNγ-treated control and mutant microglia of (G) phospho-STAT1 levels, (H) p-STAT3 levels (I) p-ERK levels and (J) p-p38 levels. n=6 for all. **p<0.01, ***p<0.001. 1-way ANOVA with Sidak’s test. (K) Volcano plots for genes differentially expressed by LPS treatment in control and TREM2 mutant microglia. (L) The number of differentially expressed genes and corrected enrichment p-value (inset) for LPS-differentially expressed genes from the ‘LPS response’ and ‘Cell activation’ GO terms. (M) Volcano plots for genes differentially expressed by IFNγ treatment in control and TREM2 mutant microglia. (N) The number of differentially expressed genes and corrected enrichment p-value (inset) for IFNγ-differentially expressed genes from the ‘Type II interferon’ and ‘Cell activation’ GO terms.

We next sought to understand the basis for this pro-inflammatory effect of the TREM2 R47H/+ mutation. We first examined expression levels of cell surface receptors for LPS and IFNγ in control and TREM2 mutant microglia. LPS bind to and activates the toll-like receptor 4 (TLR4), while IFNγ binds to the IFNγ receptor (IFNGR) heterodimer composed of IFNGR1 and IFNGR2 (Hemmi et al., 1994; Poltorak et al., 1998). However, the genes encoding these receptors were not differentially expressed in R47H/+ or T66M microglia in our RNA sequencing analysis, while qPCR analyses of microglial mRNA showed a modest downregulation of *IFNGR1* in R47H/+ and T66M cells relative to controls (Figure 3C).

We then examined the cell signaling pathways engaged downstream of LPS/TLR4 and IFNγ/IFNGR. Treatment of our microglia with LPS for 4 hours significantly increased, or trended to increase, phosphorylation of nuclear factor κB (NFκB), and the MAPKs ERK and p38 in control, R47H/+ and T66M microglia (Figures 3D-3F). While there were slight trends to an increase in the phosphorylated forms of ERK and p38 following LPS treatment in R47H/+ microglia vs. controls, these did not reach significance (Figures 3D-3F). We likewise examined induction of signaling downstream of IFNγ/IFNGR following 1-hour IFNγ treatment (Figures 3G-3J). We observed significant IFNγ-induced increases in the phosphorylation of signal transducer and activator of transcription (STAT) 1 and 3, as well as ERK and p38 MAPKs in each of control, R47H/+ and T66M microglia (Figures 3G-3J). While we again saw some very modest trends to increased phosphorylation of these proteins in R47H/+ microglia, these did not approach statistical significance (Figures 3G-3J). Thus, neither elevated receptor expression, nor hyperactive signal transduction pathways, appear likely to underlie the pro-inflammatory responses observed in TREM2 R47H/+ microglia.

Next, we characterized gene expression changes induced by LPS and IFNγ in our microglia, performing RNA sequencing on mRNA collected following two-hour treatment with these agents (Figure 3K-M). We found that LPS treatment induced similar numbers of differentially expressed genes (>0.3 log2 fold change, p-value<0.05; see Methods) in control and R47H/+ cells, while fewer genes were found altered in T66M cells (Figure 3K). As expected, differentially expressed genes from all genotypes were highly enriched for the biological process ‘response to LPS’ (Figure 3L). We noted, however, that ∼40% more genes from the ‘response to LPS’ GO category were differentially expressed in R47H/+ cells compared to controls, and that R47H/+-altered genes exhibited considerably stronger enrichment for this process than control or T66M-altered genes (Figure 3L). Furthermore, we found that LPS treatment caused differential expression of 87 genes from the ‘cell activation’ biological process in R47H/+ microglia, compared to only 50 in control and only 34 in T66M microglia, together indicating that LPS-induced gene expression changes correlate with the exaggerated cytokine release observed in microglia carrying the R47H/+ mutation.

IFNγ treatment of our microglia induced considerably more differentially expressed genes than LPS treatment (Figure 3M). ∼1450 genes were differentially expressed in R47H/+ microglia, compared to only ∼850 in control, and ∼1100 in T66M microglia, suggesting a stronger IFNγ response in R47H/+ cells (Figure 3M). Again, as expected these differentially expressed genes were strongly enriched for the pathway ‘type II interferon’, with slightly more pathway genes altered in R47H/+ microglia than control or T66M cells (Figure 3N). As with LPS, we also observed differential expression of ∼35-40% more ‘cell activation’ genes by TREM2 R47H/+ microglia following IFNγ treatment than either control or T66M cells (Figure 3N). Thus, our results indicate that in addition to a pro-inflammatory gene signature under baseline conditions (Figure 1), R47H/+ cells exhibit exaggerated gene expression changes following inflammatory insults, as well as hyper-responsive cytokine release (Figure 3).

### TREM2 R47H/+ microglia have an impaired response to laser-induced injury

In the rodent brain, microglia exhibit robust responses to various types of neuronal injury (Li and Barres, 2018). In one common injury model, microglia extend processes and migrate toward sites of laser-induced damage, a process that is impaired in heterozygous and homozygous TREM2 knockout mice (Mazaheri et al., 2017; Sayed et al., 2018). We thus sought to establish an *in vitro* laser injury response model using iPSC-derived cells to examine whether injury responses were affected by TREM2 R47H/+ or T66M mutations (Figure 4A). To that end, we generated co-cultures of wild-type Neurogenin 2 (NGN2)-induced neurons and our control and TREM2 mutant microglia (see Methods; Figures 4A and 4B). For neuronal injury, microglia were live-labeled with Vybrant DiO and then added in 25% Matrigel to overlay >3-week NGN2 cultures (Figure 4C). Under these conditions microglia concentrated around aggregates of neuronal cell bodies (Figure 4C, top panel). After 7-10 days of co-culture, neuronal injury was induced using high intensity confocal laser scanning (corresponding to the photobleached area in Figure 4C, middle panel), after which we monitored microglial responses using time lapse imaging (Figure 4C, bottom panel). We then used cell tracking software to quantify the movement of every microglia in a field of view, across time, following neuronal injury (see Methods, Figures 4C and 4D). Averaging the net movement, over 2 hours, of every tracked cell towards the center of the laser induced injury site, revealed that control, R47H/+ and T66M microglia all exhibited directed movement towards the injured area (Figure 4D). We found, however, that the injury response of R47H/+ microglia was significantly impaired, with tracked cells moving on average only half as far toward the injury site as control microglia (Figure 4D). T66M microglia also exhibited qualitatively reduced directed movement compared to control cells, though the effect did not reach statistical significance (Figure 4D).

**Figure 4.**
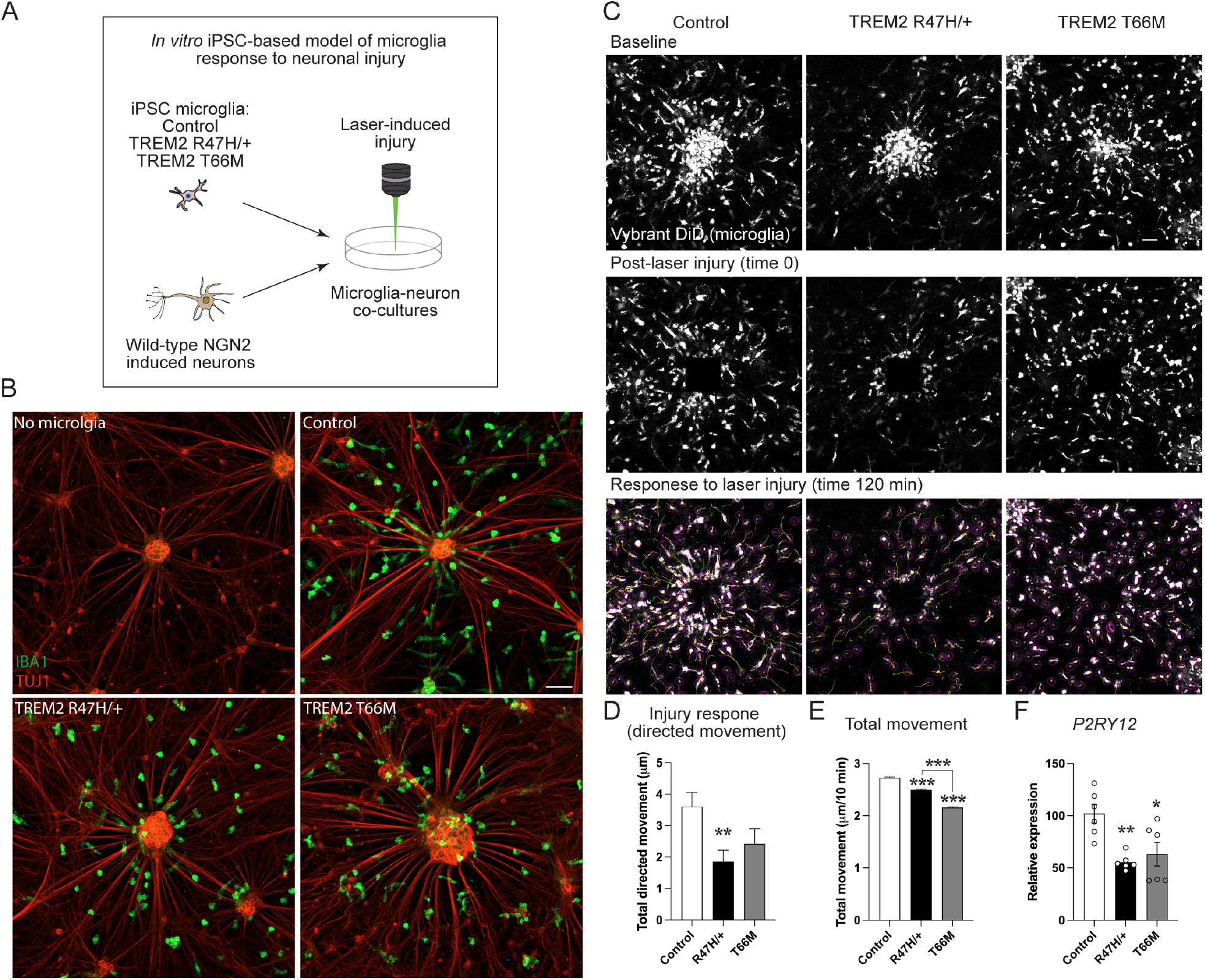
The TREM2 R47H/+ mutation impairs microglial response to laser-induced injury. (A) Schematic of laser injury experiment. (B) Images of NGN2 induced-neuron co-cultures with control and TREM2 mutant microglia stained with β-Tubulin 3 (TUJ1) (red) and IBA1 (green). Scale bar = 50 μm. (C) Images of live NGN2 induced-neuron co-cultures with control and TREM2 mutant microglia. Microglia are labeled with Vybrant DiD dye (white). Imaged before neuronal injury (top), immediately following neuron injury (middle) and 2 hours after injury induction (bottom). Identified cells (circles) and their movement tracks following injury (yellow lines) are indicated. Scale bar = 50 μm. (D) Quantification of directed movement for each cell toward the center of the injury site over the 2-hour session. >1000 cells tracked for all. n=11 time series for control (7 from clone A, 4 from clone B), n=9 time series for R47H/+ (3 from clone A, 6 from clone B), n=7 time series for T66M (3 from clone A, 4 from clone B). **p<0.01. 1-way ANOVA with Tukey’s test. (E) Quantification of total movement (in any direction) for each cell between time points. >25000 cells tracked for all, from the same time series as in (D). ***p<0.001. 1-way ANOVA with Tukey’s test. (F) Quantification of *P2RY12* mRNA levels in TREM2 mutant and isogenic control microglia. n=6 for each. *p<0.05, **p<0.01. 1-way

We next examined the total movement of microglia in these same experiments (including movement away from, and at oblique angles to, the injury site; Figure 4E). While R47H/+ and T66M microglia showed highly significant reductions in total movement compared to control cells, consistent with effects we saw in the scratch assay (Figures 2I-2K), these effects were qualitatively much less (∼9% and ∼20%, respectively) than the reductions we observed in directed movement towards injury sites (Figure 4E). Thus, it is likely that mechanisms beyond impaired cellular movement *per se*, contribute to the reduced neuronal injury response. Accordingly, we noted a downregulation of the purinergic receptor *P2RY12* in TREM2 R47H/+ microglia in our RNA sequencing analysis, as well as in both R47H/+ and T66M cells in qPCR experiments (Figure 4F). As microglial injury responses in mice are critically dependent on signaling via P2RY12, these findings suggest that impaired ATP/ADP sensing at least in part underlies the impaired injury response by R47H/+ microglia (Haynes et al., 2006).

### Xenotransplanted TREM2 R47H/+ microglia reduce synaptic density in the mouse brain

To examine our control and TREM2 mutant microglia *in vivo*, we next performed xenotransplantation experiments in immunodeficient hCSF1-expressing mice (see Methods; Hasselmann et al., 2019). Microglia were differentiated and trans-cranially injected into the hippocampi of P0-P4 pups. At ∼3 months of age, mice were sacrificed and analyzed in immunostaining experiments. Staining of transplanted brain sections for the microglia marker IBA1 and the human nuclei-specific marker STEM101, demonstrated co-localization of IBA1 and STEM101 signal in a subset of microglia (Figure 5A). Endogenous STEM101-negative mouse microglia intermingled with transplanted STEM101-positive cells, with human microglia exhibiting stronger IBA1 signal than their mouse counterparts (Figure 5A).

**Figure 5.**
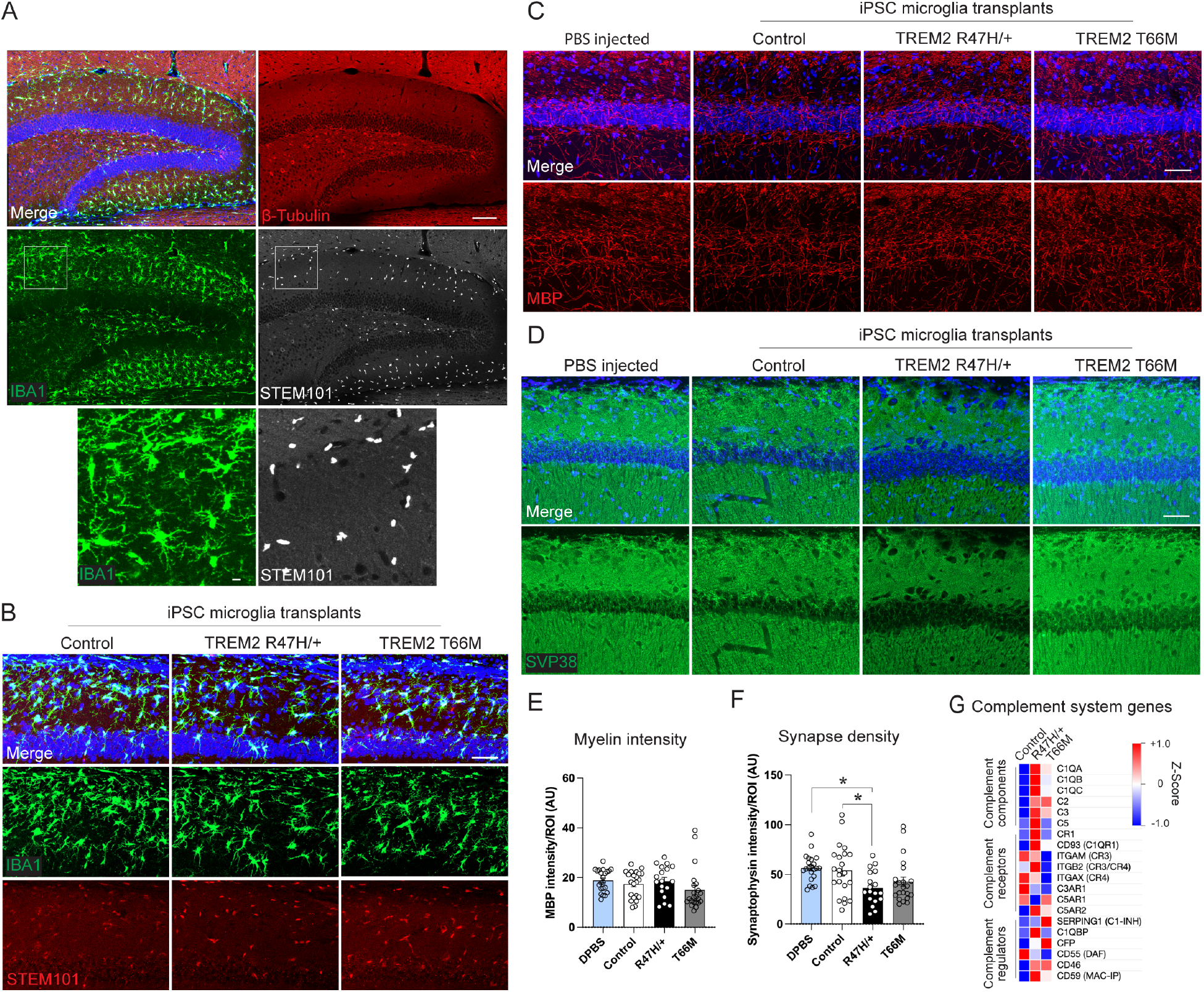
Xenotransplantation of TREM2 R47H/+ iPSC-microglia reduces synaptic density in the mouse hippocampus. (A) Tile-scan image of representative hippocampus following xenotransplant using control microglia, stained with DAPI (blue), the neuronal marker β-Tubulin 3 (TUJ1, red), the microglia marker IBA1 (green) and the human nuclei-specific marker STEM101 (white). Scale bar = 100 μm. (B) Images of transplanted control, R47H/+ and T66M microglia in hippocampal area CA1, stained for DAPI (blue), IBA1 (green) STEM101 (red). Scale bar = 50 μm. (C) Images of DAPI (blue) and MBP (red) in hippocampi from mice transplanted with PBS or control and mutant microglia. Scale bar = 50 μm. (D) Images of DAPI (blue) and SVP38 (green) in hippocampi from mice transplanted with PBS or control and mutant microglia. Scale bar = 50 μm. (E) Quantification of MBP signal in hippocampi of transplanted mice. n=22 for PBS from 11 mice, 22 for control from 11 mice, 18 for R47H/+ from 9 mice and 22 for T66M from 11 mice. (F) Quantification of SVP signal in hippocampi of transplanted mice. n=22 for PBS from 11 mice, 22 for control from 11 mice, 18 for R47H/+ from 9 mice and 22 for T66M from 11 mice. *p<0.05. 1-way ANOVA with Tukey’s test. (G) Heatmap of the gene expression Z-Score in control and TREM2 mutant microglia for complement system genes. Genes that were outside top 10,000 most expressed genes in our iPSC microglia were omitted.

We first quantified the density of transplanted control, R47H/+ and T66M microglia in area CA1 of the hippocampus, finding no significant differences in microglia numbers (Control = 29.5+/-5.4, R47H/+ = 30.4+/-3.0, T66M = 38.0+/-4.0 STEM101 cells/field of view; Figures 5B). TREM2 loss-of-function mutations in mice have been reported to impair the clearance of myelin debris in models of myelin damage, as well reducing developmental synaptic pruning, however the effects of the R47H/+ and T66M mutations have not been examined in either case (Filipello et al., 2018; Poliani et al., 2015). We also observed reduced uptake of isolated myelin and synaptosomes by R47H/+ and T66M microglia in our *in vitro* experiments as well (Figure 2). Thus, we next examined possible impacts of iPSC-microglia on myelin integrity and synapse density in the mouse hippocampus, using PBS injected mice as controls. To assess myelin integrity, we performed staining for the myelin component myelin basic protein (MBP), finding no change in signal intensity in sections from control, R47H/+ or T66M-injected mice compared to those injected with PBS (Figures 5C and 5E). Next, we assessed synapse density by staining for the synaptic protein synaptophysin (SVP38). While we found no significant change in synaptophysin intensity in brains transplanted with control or T66M microglia compared to those injected with PBS, we found a surprising and specific reduction in synaptophysin signal brains that had been transplanted with R47H/+ microglia (Figures 5D and 5F).

While we observed reduced uptake of isolated synaptosomes by R47H/+ microglia in our *in vitro* experiments, distinct mechanisms govern synapse loss *in vivo* (Hammond et al., 2019). Inflammatory insults can cause synapse loss in the mouse brain, and inappropriate activation of complement-mediated synaptic pruning has also been demonstrated in neuropsychiatric and neurodegenerative disease models (Lee et al., 2019; Rao et al., 2012). In addition to the pro-inflammatory effect that TREM2 R47H/+ mutations confer on microglia (Figures 1 and 3), our RNA sequencing experiments also indicated an upregulation of multiple complement pathway components in R47H/+ microglia (Figure 5G). In particular, C1QA, C1QB, C1QC, C3 and ITG2B (a subunit of compliment receptor 3) were all among the top 150 expressed genes in iPSC-microglia, and exhibited increased expression in TREM2 R47H/+ microglia compared to control and T66M cells (Figure 5G). Similar effects were seen for the receptors CR1 and C1QR1 in R47H/+ cells, though these genes were not as highly expressed in our microglia (Figure 5G). Thus, these observations indicate that TREM2 R47H/+ microglia can promote synapse loss *in vivo*, likely mediated by inflammatory mechanisms and/or an activation of complement-mediated synaptic pruning.

## Discussion

Taken together, our observations indicate shared and distinct effects of the TREM2 R47H/+ and T66M mutations on microglial gene expression and function. We identified a core set of genes that were differentially expressed by both mutations, affecting vesicle mediated transport, lipid metabolic processes, apoptosis, and cellular movement among other processes. We also found similar functional impairments in cellular movement and the uptake of multiple brain-relevant substrates in cells carrying the R47H/+ and T66M mutations.

Importantly, our findings demonstrate that the TREM2 R47H/+ mutation has many additional effects on microglia that are not seen in cells carrying the T66M mutation. A large set of genes were specifically altered in R47H/+ microglia, most notably affecting immune and inflammatory pathways and cell activation. Genes regulating phosphorylation and phosphate metabolic pathways, cytoskeletal organization and the complement system were also altered by the R47H/+ mutation. These unique gene expression changes were mirrored at the functional level, with R47H/+ microglia exhibiting enhanced cytokine release and gene expression changes following inflammatory insult, and only R47H/+ cells showed a significant deficit in laser injury response. We also observed that TREM2 R47H/+ microglia induced synapse loss when transplanted into mouse brains. While there is still not consensus in the field, our findings strongly support a growing body of evidence that TREM2 R47H/+ mutations do not act as simple loss-of-function mutations (Cheng-Hathaway et al., 2018; Liu et al., 2020; Piers et al., 2020; Sayed et al., 2021; Song et al., 2018).

Chronic neuroinflammation is a well-known feature of neurodegenerative disease that can result from protein aggregates and neuronal damage among many other causes (De Strooper and Karran, 2016). Importantly, genetic factors such as APOE4, the most common cause of AD, have also been shown to modulate intrinsic inflammatory responses by brain cells (Ennerfelt and Lukens, 2020; Lin et al., 2018; Vitek et al., 2009). While previous studies in mice and human iPSC models have reported both pro- and anti-inflammatory phenotypes associated with R47H mutations, our observations support recent snRNA sequencing and functional studies showing a pro-inflammatory effect of TREM2 R47H/+ mutations (Liu et al., 2020; Sayed et al., 2021).

Synapse loss is another key event in AD progression. It is the pathological feature that correlates most closely with cognitive decline, and often precedes neuronal loss (Canter et al., 2016). Inflammatory insults can promote the loss of synaptic elements, while complement-mediated synaptic pruning can also become inappropriately activated in a number of neurological conditions (Hammond et al., 2019; Hong et al., 2016; Lee et al., 2019; Rao et al., 2012). Indeed, the complement receptor 1 (CR1) is a risk factor for AD, while polymorphisms affecting complement components C3 and C4 are associated with schizophrenia risk (Lambert et al., 2009; Lee et al., 2019; Sekar et al., 2016). While TREM2 knockout mice exhibit impaired developmental synaptic pruning, our transplantation experiments show that microglia carrying the TREM2 R47H/+ mutation can promote synapse loss in the mouse brain (Filipello et al., 2018). In addition to a pro-inflammatory phenotype that could impact synapse integrity, we found that R47H/+ microglia exhibited up-regulated expression of multiple complement pathway genes, including the AD risk factor CR1 and the schizophrenia risk factor C3. Notably, the complement genes C1QA, C1QB, C1QC and C3 were among the most highly expressed genes in our iPSC-microglia, and were further upregulated by the R47H/+ mutation. Furthermore, C1QC and C3 were among the genes found upregulated in microglia from the brains of AD patients carrying the R47H/+ mutation compared to those with wild-type TREM2, underscoring the likely disease relevance of our findings (Sayed et al., 2021).

While our observations show gene expression and functional alterations in microglia carrying TREM2 T66M mutations, most were also observed on R47H/+ cells. Thus, it remains unclear why T66M mutations in patients result in more severe and more highly penetrant neurodegenerative disease than do R47H/+ mutations. One possibility is the enrichment in fibrinolysis regulators we observed in T66M upregulated genes, none of which were differentially expressed among shared, or R47H/+-specific altered genes. In addition to their roles in blood clotting, fibrinolysis regulators can modulate extracellular matrix remodeling and blood-brain barrier permeability that can impact brain health (Patel et al., 2008; Ziliotto et al., 2021). Blood brain barrier damage and the presence of auto-antibodies have been reported in PLOSL patients, potentially due to dysregulated fibrinolysis (Errichiello et al., 2019; Kalimo et al., 1994). Additional studies, however, will be required to establish the exact functions of fibrinolysis regulators, and of TREM2 T66M mutations more generally, in the etiology of neurodegenerative disease.

Together, these findings improve our understanding of TREM2 biology, and uncover numerous gene expression and functional effects that arise from the TREM2 R47H/+ and T66M mutations in human microglia. While identifying a core set of genes and cellular functions that are altered by both TREM2 R47H/+ and T66M mutations, we also find multiple processes that are uniquely affected by the R47H/+ AD variant. Prime among these are a strong pro-inflammatory phenotype, and an activation of complement pathway components, both of which may contribute to microglial R47H/+-mediated synapse loss.

Thus, our findings highlight multiple effects of the TREM2 R47H/+ mutation likely to underlie its association with AD risk, as well as suggesting new nodes that could be exploited for therapeutic intervention.

## Acknowledgements

We thank and P.C. Pao for thoughtful comments on the manuscript; E. McNamara for mouse colony maintenance and related support; Y. Zhou, M. Mazzanti, and T. Garvey for administrative support. Our work in the Tsai lab is only possible through the generous support of The Robert A. and Renee E. Belfer Family Foundation. This work was also supported by a Long-Term Fellowship postdoctoral award to J.P. from the Human Frontier Science Program.

## Methods

### Gene editing, induced pluripotent stem cell and microglia culture

iPSC culture was performed in the MIT Picower Institute iPSC core facility. iPSCs from an unaffected 75-year-old female were used as the parental line (Coriell #AG19173). Cells were cultured on hES-Qualified MatrigelR (Corning)-coated tissue culture plates in mTeSR1 media (STEMCELL). CRISPR-Cas9-mediated gene editing to generate TREM2 R47H/+ and T66M mutations was performed as previously described (Lin et al., 2018). Potential edited clones were screened by Sanger sequencing (Azenta Life Sciences), after which we selected two clones carrying each of R47H/+ or T66M-causing mutations for further analysis. An additional clone from the CRISPR screening that remained un-edited at the TREM2 locus, as well as the parental iPSC line, were used as isogenic controls. Cells were karyotyped to identify any chromosomal abnormalities and the top 4 predicted CRISPR off-target sites were sequenced to rule out off-target mutations. Differentiation to iPSC-microglia was performed according to McQuade et al. (2018) with modified maturation supplementation using 25 ng/mL hM-CSF and 100 ng/mL hIL-34 (PeproTech) (McQuade et al., 2018). *In vitro* experiments were performed using DIV 40-60 microglia.

### Quantitative PCR

RNA was extracted from microglia using the QIAGEN RNeasy Plus Mini Kit. cDNA synthesis was performed with RNA to cDNA EcoDry Premix (Oligo dT) (Clontech). qPCR was performed with SsoFast EvaGreen Supermix (Bio-Rad) using a C1000 Thermal Cycler and a C96 Real-Time System (Bio-Rad). Target genes were normalized using β-Actin (ACTB).

### Western blotting

Protein was extracted using RIPA buffer. Western blots were performed using PVDF membranes (Millipore) following standard methods. Antibodies used included: goat-TREM2 (Cell Signaling Technology), mouse-β-actin (Sigma), rabbit-p-AKT (S473, Cell Signaling Technology), mouse-AKT (Cell Signaling Technology), rabbit-pan-p-PKC (T514, Cell Signaling Technology), rabbit-p-ERK (Cell Signaling Technology), rabbit-ERK (Cell Signaling Technology), rabbit-p-p38 (T180/182, Cell Signaling Technology), rabbit-p38 (Cell Signaling Technology), rabbit-p-NFkB (S536, Cell Signaling Technology), mouse-NFkB (Cell Signaling Technology), rabbit-p-STAT1 (T701, Cell Signaling Technology), rabbit-STAT1 (Cell Signaling Technology), rabbit-p-STAT3 (T705, Cell Signaling Technology), rabbit-STAT3 (Cell Signaling Technology).

### RNA Sequencing and analysis

RNA was extracted from microglia using the QIAGEN RNeasy Plus Mini Kit and subject to QC using an Advanced Analytical-fragment Analyzer before library preparation using NEB Ultra II RNAseq library preparation kit. Libraries were pooled for sequencing using Illumina NextSeq500 or NovaSeq6000 platforms at the MIT BioMicro Center. Reads were aligned to the human genome reference GRCh38/hg38 using the R software package Rsubread (Liao et al., 2019). Mapped reads were converted to gene level counts using the featureCounts function of Rsubread with “strandSpecific” parameter set to 0. Differential analysis was performed using DESeq2 (Love et al., 2014). Differential expression for inflammatory treatments was performed comparing baseline, LPS and IFNγ within genotypes. Cutoffs for differentially expressed genes in our initial analysis of control, R47H/+ and T66M microglia were set at >0.3 log2 fold change, p-adjusted <0.05, among the top 5000 expressed genes in our iPSC-microglia. ToppGene was used for gene ontology analysis (Chen et al., 2009). For LPS and IFNγ-induced gene analysis, cutoffs were set at >0.3 log2 fold change, p-value <0.05. Here the top 10,000 expressed genes in iPSC-microglia were used as many cytokine-genes are expressed at low levels without induction. ToppGene was again used for gene ontology analysis. Morpheus (Broad Institute, https://software.broadinstitute.org/morpheus) was used to generate heatmaps of the Z-Score for expression of inflammatory and complement genes across genotypes.

### Uptake, immunostaining, imaging and flow cytometry

Synaptosomes were isolated as previously described (Penney et al., 2017). Myelin was isolated using a modified protocol from Larocca and Norton (Larocca and Norton, 2006). Briefly, wild-type mouse brains were homogenized in 0.32 M sucrose with a loose pestle. Homogenate was layered on top of 0.85 M sucrose and spun at 42,865 g for 60 min. The interphase was added to ultrapure water (Thermo Fisher) and spun at 42,865 g for 15 minutes. The pellet was then washed twice in ultrapure water (6915 g, 15 minutes), resuspended in 0.32 M sucrose, layered on top of 0.85 M sucrose and spun at 42,865 g for 30 minutes. The interphase was subsequently transferred and washed in 10 mL ultrapure water (42,865 g, 15 minutes). Amyloid-β_42_-HiLyte Fluor 488 was obtained from AnaSpec. Uptake for immunostaining experiments was performed in 24-well plates containing glass coverslips. 20 μg/mL synaptosomes, 40 μg/mL myelin or 200 ng/mL Aβ_42_-488 was added to culture media. 3 hours later cultures were washed and fixed with 4% paraformaldehyde (Electron Microscopy Solutions) in phosphate-buffered saline (PBS; Thermo Fisher) for 10 minutes. After 3X PBS washes, cells were blocked 1 hour with 5% normal goat serum (Millipore) and 0.2% Triton-X 100 (Sigma). Staining was performed overnight at 4°C. Neuron-microglia co-cultures were fixed similarly; co-cultures and xenotransplant brain sections were stained using the same protocol. Antibodies used included: mouse-SVP38 (Sigma), chicken-MBP (Millipore), guinea pig-IBA1 (Synaptic Systems), mouse-β-Tubulin 3 (TUJ1; GeneTex), rabbit-β-Tubulin 3 (BioLegend) and mouse-STEM101 (Takara Bio). Imaging was performed using Zeiss LSM880 or LSM900 confocal microscopes. For flow cytometry, synaptosomes or myelin were conjugated with pHrodo iFL Green STP Ester (Invitrogen) at a 1:10 pHrodo:synaptosome/myelin ratio, incubated 30 minutes, pelleted, then washed 2X with PBS, pelleting each time. 20 μg/mL pHrodo-synaptosomes, 40 μg/mL pHrodo-myelin or 200 ng/mL Aβ_42_-488 was added the media of microglia plated in 96-well plates (20,000/well) and incubated 3 hours. Flow cytometry was performed using a FACS Celesta HTS-1 sorter (Koch Institute Flow Cytometry Core at MIT).

### Scratch assay

Microglia were plated at 400k per well in 6-well plates. The next day, cells were physically removed (scratched) from the middle portion of the well, and media was changed to remove floating cells. Images were collected immediately after scratch, and 2 days later using an EVOS FL microscope (Thermo Fisher) to monitor movement back into the scratch area.

### Inflammatory treatments and ELISA

ELISAs were performed using culture media from control and TREM2 mutant microglia grown in 96-well plates (20k/well). Soluble TREM2 was quantified using the human TREM2 ELISA Kit (Abcam). For IL6 and TNFα ELISAs, cells were either left untreated, or treated for 2 days with 100ng/mL LPS or 50 ng/mL IFNγ before collection of culture media. Human IL-6 or TNF-alpha Quantkine ELISA Kits (R+D Systems) were used. For RNA sequencing experiments, 200k microglia in 12-well plates were treated with 100ng/mL LPS or 50 ng/mL IFNγ for 2 hours before RNA extraction.

### Live imaging

Live-imagining was performed using a Zeiss LSM900 equipped with a humidity and CO_2_ controlled, heated chamber kept at 37°C. For live uptake experiments, microglia were labeled with Vybrant DiD (Molecular Probes) and plated at 200k/well in 12-well plates. 200 ng/mL Aβ_42_-488 was added to the media and images were collected every 10 minutes for 2 hours. Aβ_42_-488 signal co-localizing with Vybrant DiD-labeled microglia was quantified across time to monitor uptake.

### In vitro laser injury model

NGN2-induced neurons were initially grown on Matrigel-coated plates, then transferred to poly-D-lysine (Sigma-Aldrich) coated plates at day 7 of induction, at which point they were grown in BrainPhys Neuronal Media (STEMCELL) supplemented with 2% B27 (Life Technologies), 1% N2 (Life Technologies), 1% NEAA (Sigma-Aldrich). 1% GlutaMAX (Thermo Fisher), 200 nM L-Ascorbic acid, 20 ng/mL hBDNF (PeproTech), 20 ng/mL hGDNF (PeproTech). Neuron-microglia co-cultures were grown in the same media but supplemented additionally with 25 ng/mL hM-CSF. DIV21 or older NGN2 cultures (∼250k cells/24-well plate) were overlaid with ∼250k Vybrant DiD (Molecular Probes)-labeled iPSC-microglia in 200 μl 25% Matrigel:75% BrainPhys media. After 30 minutes, warm BrainPhys media (without Matrigel) was returned to the cultures. Following 7-10 days of co-culture, Vybrant DiD microglia were live-imaged on a Zeiss LSM900 with a temperature/CO_2_ controlled chamber to capture baseline state. Laser injury was induced at 6X zoom by 3 minutes live scanning at 100% laser intensity, focused to the middle of a neuron cell body cluster. A photobleached area, corresponding to the laser-induced injury, was evident following laser scanning injury. Images of Vybrant DiD microglia signal were captured every 5 minutes post-injury for 2 hours. Microglia movement in the image series was tracked using the Fiji TrackMate plugin (Tinevez et al., 2017). The vectors for x-y displacement of each tracked cell between each time point were used to determine total microglial movement. Directed movement towards (or away from) the injury site was calculated as the difference between the initial and final distance of a given cell from the center of the injury site, for each cell that could be tracked for the entire 2-hour session.

### Xenotransplant experiments

Rag2 knockout, IL-2rγ knockout, human CSF1-expressing mice were housed in the MIT Department of Comparative Medicine SCID facility. Transplants were performed as described by Hasselmann and colleagues (Hasselmann et al., 2019). Briefly, following HPC induction, ∼500k microglia precursors were injected transcranially into early postnatal (P0-P4) Rag2 knockout, IL-2rγ knockout, human CSF1 pups using a Hamilton syringe. At 3-4 months of age, transplanted mice were anaesthetized, perfused with PBS, then with 4% paraformaldehyde in PBS before drop-fixing brains in 4% paraformaldehyde overnight. After washing with PBS, brains were sectioned at 40 μm thickness on a Leica VT1000S vibratome. Brain sections were then immunostained and imaged as described in the ‘*Uptake, immunostaining, imaging and flow cytometry*’ section above.

### Statistical analysis

Data are presented as mean +/-SEM and were analyzed using Prism 9 (GraphPad). 1-way ANOVA followed by post hoc Tukey’s test was used for most analyses, with Sidak’s multiple comparison test used to compare baseline and LPS/IFNγ-induced conditions within genotype, as well as LPS/IFNγ-induced conditions between genotypes. In all cases *p<0.05 was considered significant.

